# Riceberry rice bran protein hydrolyzed fractions induced apoptosis, senescence and G1/S cell cycle arrest in human colon cancer cell lines

**DOI:** 10.1101/2021.08.20.457150

**Authors:** Vichugorn Wattayagorn, Mesayamas Kongsema, Sukuntaros Tadakittisarn, Pramote Chumnanpuen

## Abstract

Riceberry rice bran is the part of rice that has been scrubbed out during coloring process. There are various health benefits with high protein content and antioxidant ability. The hydrolyzed rice bran consists of diverse peptides that provide various bioactive properties. This work aimed to study the effect of hydrolyzed riceberry rice bran extracted on colon cancer cell lines (HT-29 and SW-620) compared to normal cell (PCS-291-010). The MTT assay result showed that our extract has less cytotoxicity on normal cell (PCS-291-010, IC_50_ = 6,680.00 μg/ml) compared to the colon cancer cell lines and has more effect on metastatic cancer cell line (SW-620, IC_50_ = 5,492.31 μg /ml) than non-metastatic cancer cell line (HT-29, IC_50_ =6,040.76 μg/ml). According to the DNA fragmentation pattern analysis, the ladder pattern indicated that the rice bran extract can induce the apoptosis process in SW-620 cell line. Confirmed the pattern of apoptotic cell by AO/PI double stain test and quantified apoptotic cells by Annexin V. For the cell senescence analysis, SA-β-gal staining technique was performed at 24 h after treatments, HT-29 reached maximum senescence rate at 85.74% while SW-620 had only 17.23% of senescence. And a result of cell cycle analysis, HT-29 were decreased the number of cells in S, M/G2 phase, and increased the number of cells in G0/G1 phase. Furthermore > 50 kDa peptide fraction separated from HRBE has a potent anti-cancer cells (SW-620, IC_50_ = 4,908 μg/ml). In conclusion, the hydrolyzed riceberry rice bran extract can inhibit colon cancer cell lines with less effect on normal cell. The extracts could induce apoptosis process in metastatic cancer cell and induce senescence process in non-metastatic cancer cell. This observed information will be useful and applicable for medical research and colon cancer treatment in the future.

## Introduction

Colon cancer is a major health problem globally and one of the three most common cancer diagnosed in both men and women worldwide. Its incidence rapidly rises in cities with economic development. Colon cancer can be caused by accumulating mutations and epigenetic alterations [1]. Presently, various therapies have been used clinically for colon cancer, including chemotherapy. However, chemotherapy resistance remains one of the serious problems and can also cause adverse side effects. Thus, the development of anti-cancer natural extracts could be possibly the alternative treatment for future colon cancer patients to overcome those problems.

Rice (Oryza sativa L.) is the most important crop and the staple food source con-sumed by over half of the world population. Thailand is the highest production for ag-riculture. Recently, riceberry rice has become popular among health-concern people due to the health-promoting and potential anti-cancer activity. Moreover, Riceberry rice has also been reported to have hypoglycemic, hypolipidemic, antioxidant, and an-ti-inflammation properties. The active ingredients of riceberry were tocopherol, tocotrienol, β-Sitosterol, γ-oryzanol, β-Carotene and some protein. These substances were found in most of pericarp, also known as rice bran [2]. From previous studies, riceberry rice bran extract was significantly inhibited on colonic carcinoma (Caco-2), breast adenocarcinoma (MCF-7) and acute myeloid leukemia (HL-60) by suppressing cell growth and proliferation, inducing cell cycle arrest and apoptosis [3]. However, the use of protein hydrolysate from riceberry rice bran on colon cancer inhibition remains limited. Although the rice bran was rich of protein such as albumin, globulin, glutelin and prolamin with high nutritional value, minerals, unsatured fat and vitamins. And highest phytate content was found in albumin and the lowest in prolamin [4–6] that can hydrolysates by Alcalase enzyme. According to a previous study, Alcalase is an alkaline endo-peptidase that gave higher protein yields. Its activity in hydrolyzing glutelin (MW 57 kDa), as the main protein accounting for 60-80% of total protein in rice [7, 8].

Thus, the purpose of this study is to investigate the anti-cancer activity of hydrolyzed riceberry bran extract on non-metastatic and metastatic colon cancer cell lines. This investigation may give useful information which leads to the future application in medical research and treatment for colon cancer patients.

## Materials and Methods

### 1. Riceberry rice bran hydrolysation, extraction and fractionation

Hydrolyzed riceberry rice bran by alcalase was performed according to Thamnarathip et al., 2016[26]. The hydrolyzed riceberry rice bran powder was mixed with ultrapure water in a concentration of 20 g/L. Then, the mixture was centrifuged at rpm 3,600 degrees for 5 minutes. Filter with whatman filter paper No. 1, size 11 μm before Freeze-drying process. The freeze-dried powder is then mixed with ultrapure water at a concentration of 50 mg/ml before being filtered with 0.2 μm syringe filter to sterile the mixture. Hydrolyzed riceberry rice bran extract (HRBE) mixture will be kept at −20 °C until further use. Then, isolated size of peptide by Amicon Ultra-15 centrifugal filter devices into >50, 50-30, 30-10, 10-3 kDa, and <3 kDa fractions. And freeze-drying before test cytotoxicity with SW-620 cell lines.

### 2. Extraction and isolation of anthocyanins method and fractionation

Isolation of anthocyanins from HRBE to obtain pure protein by water, methanol with 0.01% HCl and acetone. The extraction method follow a study of [13]. Remove methanol and acetone in a rotary evaporator at 40°C under vacuum before Freeze-drying process. Then, isolated size of peptide by Amicon Ultra-15 centrifugal filter devices into >50, 50-30, 30-10, 10-3 kDa, and <3 kDa fractions. And freeze-drying before test cytotoxicity with SW-620 cell lines. A yield percentage was calculated by:

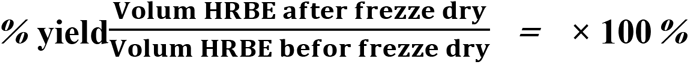

### 3. Cell culture

Fibroblast Normal Human cell line (PCS-291-010) and Non – metastasis human colon cancer cell line (HT-29) were cultured in Dulbecco’s modified Eagle’s medium (DMEM: Gibco, Thermo Fisher, USA) and metastasis human colon cancer cell line (SW-620) was cultured in Roswell Park Memorial Institute medium 1640 (RPMI 1640: Gibco, Thermo Fisher, USA). All media were supplemented with 10% fetal bovine serum (FBS) and 1% Penicillin/Streptomycin (Gibco, Thermo Fisher, USA). Cells were maintained at 37 °C in an incubator (Binder, Germany) under 5% CO_2_ and 95% air atmosphere at constant humidity. Trypsin-EDTA was used for cells detachment at 70% confluence.

### 4. Cellular viability by MTT assay

The cytotoxicity of the HRBE was determined using the MTT assay. HT-29, SW-620 and HDFa cells line were seed at 5×10^4^ cells/ml in 96-well plates. The next day, cells were treated with different concentrations of by HRBE concentrations 0 – 10,000 μg/ml and mitomycin C concentrations 0 – 100 μg/ml for 24, 48 and 72 h. To harvest, MTT solution is added and incubated for 4 h at 37 °C for the formazan. Next, the excess of MTT solution was carefully removed and replaced by 100 μl DMSO to dissolve formazan crystals. Absorbance was then measured using a microplate reader at a wavelength of 570 nm (Biotek, USA) [27]. The percentage of cell viability was quantify using following formula.

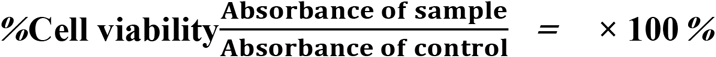

And Calculate the half-maximal inhibitory concentration (IC_50_). As HRBE demonstrated the lower IC_50_ value in SW-620 cell line, we used only SW-620 to confirm cell apoptosis in the next assay.

### 5. Senescence-associated beta-galactosidase (SA-β-gal) staining

SA-β-gal staining was performed using Senescence β-Galactosidase Staining kit (Cell signaling technology, USA) according to the manufacturer’s instruction. Briefly, the cells were examined under the inverted light microscope to monitor the morphological changes. SW-620 and HT-29 cell lines were seeded at 1×10^5^ cells/ml. in 6-well plates before treated with different concentrations of HRBE (5, 2.5 and 1 mg/ml) 0.05 mg/ml of mitomycin c for 24 h. The next day, the cells were washed three times with PBS and fixed with 4% paraformaldehyde for 15 min at room temperature. The fixed cells then were incubated overnight at 37 °C with the working solution containing 0.05 mg/mL X-gal using SA-β-gal staining kit.

### 6. The DNA fragmentation pattern analysis by gel electrophoresis technique

Agarose gel electrophoresis was the most effective way of separating DNA fragments pattern when cell apoptosis [28]. Sw-620 cells line were seeded at 1×10^5^ cells/ml in 10 cm petri dish. The next day, cells were treated with different concentrations of HRBE (10, 5 and 2.5 mg/ml) and 0.05 mg/ml of mitomycin c. The cells were extracted by DNA extraction kit and the DNA concentration was measured by Nanodrop spectrophotometer. DNA fragmentation was detected by electrophoresis using 1.5% agarose gel at a 100volte voltage for 1 hour. After that, examined under ultraviolet light with a gel documentation (Bio - Rad, UK).

### 7. Acridine orange/Propidium Iodide (AO/PI) double staining test

Acridine orange (AO) and propidium iodide (PI) were among the most used fluorescent dyes used to analyze cell culture viability and specificity for living, apoptotic and late apoptosis/necrosis states [29]. All cell lines were seed at 1×10^4^ cells/ml. in 6-well plates before being treat with different concentrations of of HRBE (10, 5, 2.5 and 1.25 mg/ml) and 0.05 mg/ml of mitomycin c for 72 h. Trypsin-EDTA was use for cells detachment and the pellet were collect by centrifugation at 1,500 rpm for 100 min, then discard the culture medium. Wash tree time with PBS and add 10 μL of PBS. Separately, 1 μL AO (10 mg/mL) was mix with 1 μL PI (1 mg/mL) making a double-staining dye of AO/PI and keep in an ice bath in dark conditions. An aliquot of the cell suspensions (10 μL) was add to the dye mixture and transferred onto a glass slide for fluorescence viewing. Slides were observed under a UV-Fluorescence microscope within 30 min.

### 8. Apoptosis assay

Apoptosis assay was perform using Muse™ Annexin V & Dead Cell Assay [30]. SW-620 cells lines were seed at 1×10^5^ cells/ml. in 6-well plates and treat with different concentrations of HRBE (10, 5, 2.5 and 1.25 mg/ml) and 0.05 mg/ml of mitomycin C for 72 h. To harvest, cells were trypsinization and collect by centrifugation at 1,500 rpm for 5 min, then wash by PBS. Cells were then move to 1.5ml eppendorf and resuspend with 50 μl complete media. Then, cells were stain with 50 μl Muse Annexin V & Dead Cell Kit (Muse, USA).and incubated at room temperature in the darkness for 20 min. The stained cells were measured using Guava Muse Cell Analyzer (Millipore, USA).

### 9. Cell cycle analysis

HT-29 cells lines were seed at 5×10^5^ cells/ml. in 6-well plates and treat with different concentrations of HRBE (10, 5, 2.5 and 1.25 mg/ml) and Positive control used 0.05 mg/ml of mitomycin C and 1 μM of Doxorubicin for 24 h. To harvest, cells were trypsinization and collect by centrifuge at 1,500 rpm for 5 min, then wash by PBS. And fix in 200 μl of ice cold 70% ethanol and incubate for at least 3 hr at – 20°C. Centrifuge at 1,500 rpm for 5 min and wash once with PBS. Then, cells were stain with 200 μl Muse Cell Cycle reagent (Muse, USA).and incubated at room temperature in the dark for 30 min. The stained cells were measured using Guava Muse Cell Analyzer (Millipore, USA).

### 10. Statical analysis

Statistical analysis for comparisons among multiple groups (3 type of cell line, Extract Concentrations and time point) were perform using analysis of variance (ANOVA) and student’s t – test. Results were expressed as mean ± standard deviation (SD). Differences with P values of < 0.05 were considered significant.

## Results

### 1. Cytotoxicity test by MTT assay

After extracting the HRBE by water got a percentage yield equal to 85.94 %. The cytotoxicity test on fibroblast normal human cell line (PCS-291-010), Non – metastasis human colon cancer cell line (HT-29) and metastasis human colon cancer cell line (SW-620) with hydrolyzed riceberry rice bran extract (HRBE) concentrations 0 – 10,000 μg/ml and mitomycin C concentrations 0 – 100 μg/ml for 24 h, 48 h and 72 h by using MTT assay.

In cancer cell lines after treated with HRBE showed a significant decreasing in cell viability with a dose and time dependent manner. The half maximal inhibitory con-centration (IC_50_) of HT-29 as 25650, 11004.50, 6040.76 μg/ml and SW-620 as 10617.19, 6767.11, 5492.31 μg/ml at 24 h, 48 h and 72 h respectively. For the effect of HRBE on Fibroblast Normal Human cell line (PCS-291-010) the IC_50_ as 19698.75, 16759.17, 16759.17 μg/ml (Fig1). For Positive control used mitomycin C on HT-29, SW-620 and PCS-291-010. The IC_50_ of HT-29 as 47183.33, 11578, 4795.41 μg/ml, SW-620 as 14487.41, 5578.10, 3850.00 μg/ml and PCS-291-010 as 19698.75, 16759.17, 16759.17 μg/ml (Fig 2) at 24 h, 48 h and 72 h respectively. The result show that the effect of HRBE has less effect on normal cells than 2 type cancer cells and the IC_50_ were decrees in dose-time dependent. The lower of IC_50_ was to tested with SW-620 cell line at 72h therefore we chose this type of cell to determine the effectiveness of the extract if it can induce apoptosis in the next experiment.

**Fig1.**
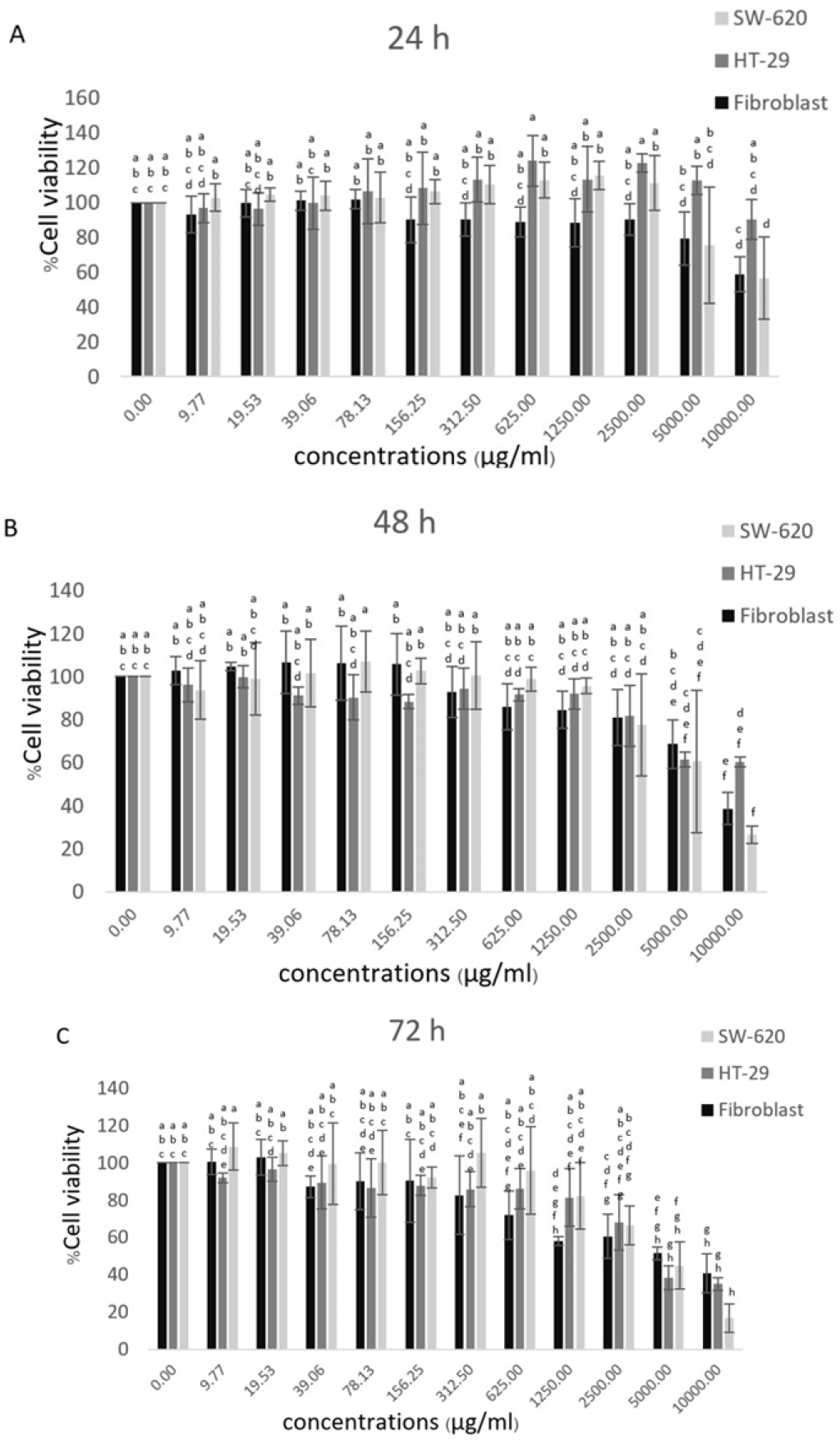
Cytotoxicity test by MTT assay on fibroblast normal human cell line (PCS-291-010), non-metastatic colon cancer cell line(HT-29) and metastatic colon cancer cell line (SW-620) after treated with HRBE concentrations (0 – 10,000 μg/ml). For 24 h (A), 48 h (B)and 72 h (C). The percentage of cell viability was determined as compared to an untreated condition. Data represented in the mean ± SD. Statistical analysis was performed using two-way ANOVA.

**Fig 2.**
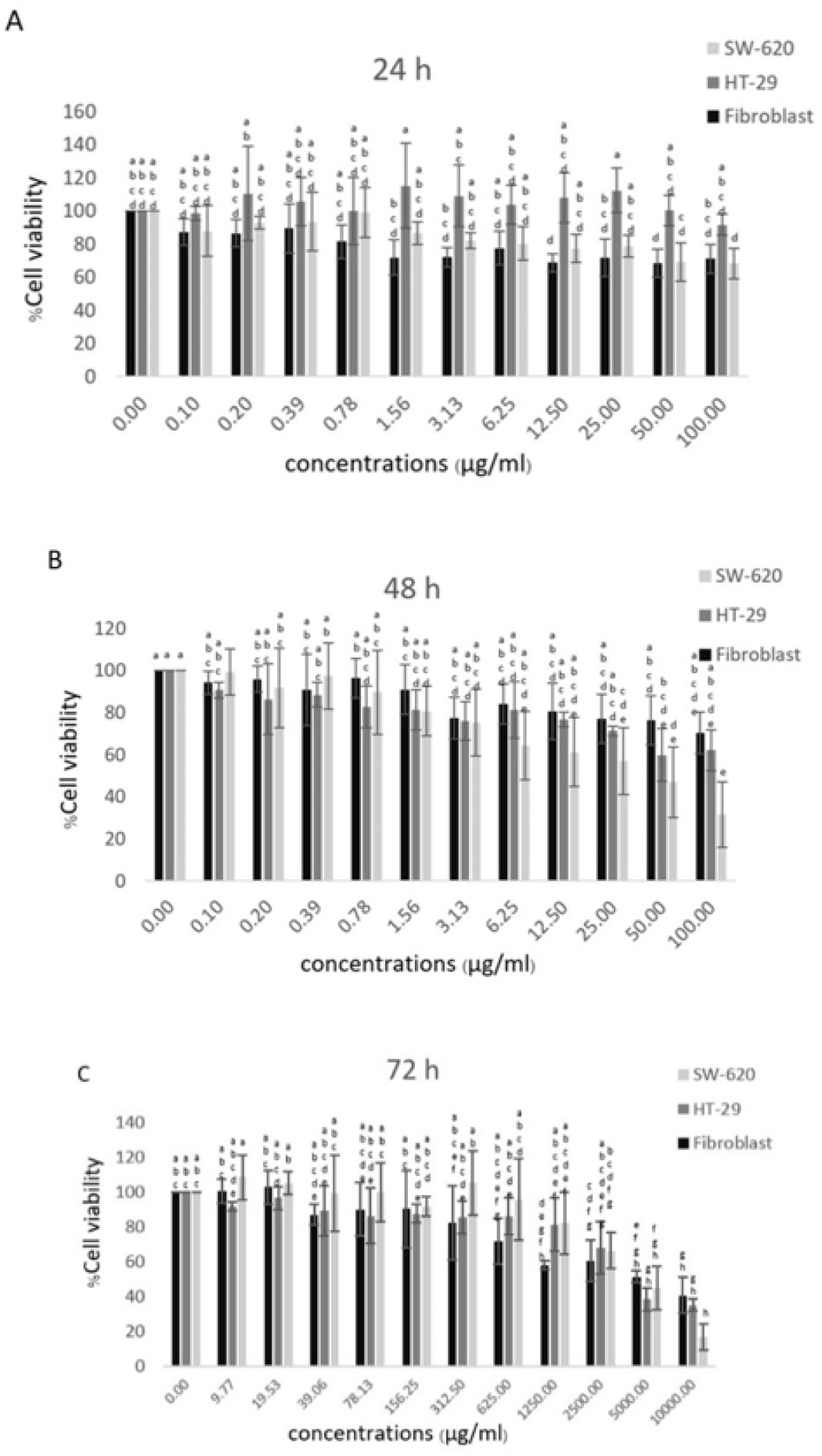
Cytotoxicity test by MTT assay on fibroblast normal human cell line (PCS-291-010), non-metastatic colon cancer cell line(HT-29) and metastatic colon cancer cell line (SW-620) after treated with mitomycin C concentrations (0 – 100 μg/ml). For 24 h (A), 48 h (B)and 72 h (C). The percentage of cell viability was determined as compared to an untreated condition. Data represented in the mean ± SD. Statistical analysis was performed using two-way ANOVA.

### 2. Apoptosis cell death by Gel-electrophoresis technique

The DNA fragmentation is being used as a marker for identification of apoptotic cells. As shown in Fig. 3, marked DNA fragmentation was observed in SW-620 cells treated with HRBE concentrations 10, 5, 2.5, 0 mg/ml and mitomycin C for 72 h. The result show the DNA bands of cells that received HRBE increase the intensity of DNA bands according to the concentration of the extracts that demonstrated cell apoptosis.

**Fig 3.**
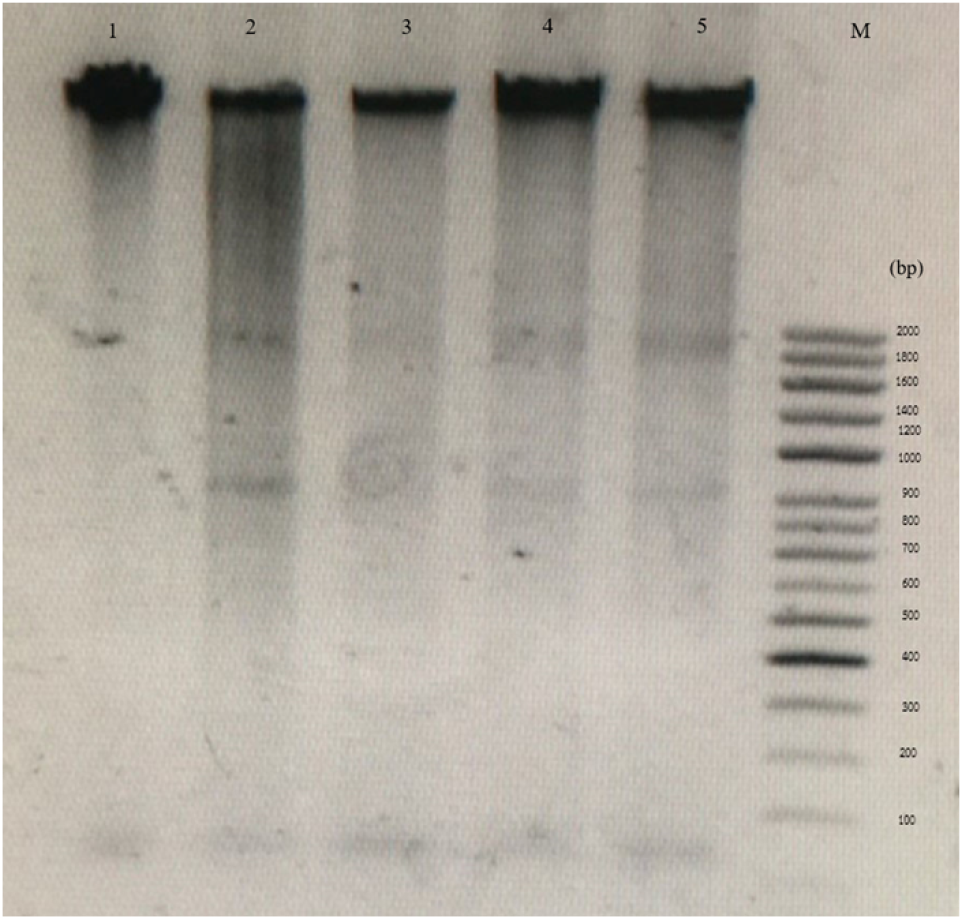
Gel electrophoresis showing DNA fragmentation pattern of metastatic colon cancer cell line (SW-620). After treated with mitomycin C and HRBE at different concentrations for 72 h. Standard DNA marker 100 bp DNA ladder geneSTA for DNA fragment size estimation (bp)

Lane 1: control
Lane 2: 0.05 mg/ml mitomycin C
Lane 3: 2.5 mg/ml HRBE
Lane 4: 5.0 mg/ml HRBE
Lane 5: 10.0 mg/ml HRBE
Lane 6: standard DNA marker

### 3. Senescence-associated beta-galactosidase (SA-β-gal) staining

Cell senescence have noticeable morphological changes in morphology, such as enlarged and flattened cell shape and increased granularity and biochemical changes. The most common assay used to check cellular senescence is Senescence-associated beta-galactosidase (SA-β-gal) staining. In this study of the ability of HRBE can induced senescence in HT-29 and SW-620 cells at concentrations of 0, 0.5, 1, 2.5, 5 mg /ml and mitomycin C 0.05 mg / ml (Figs 4 and 6).

**Fig 4.**
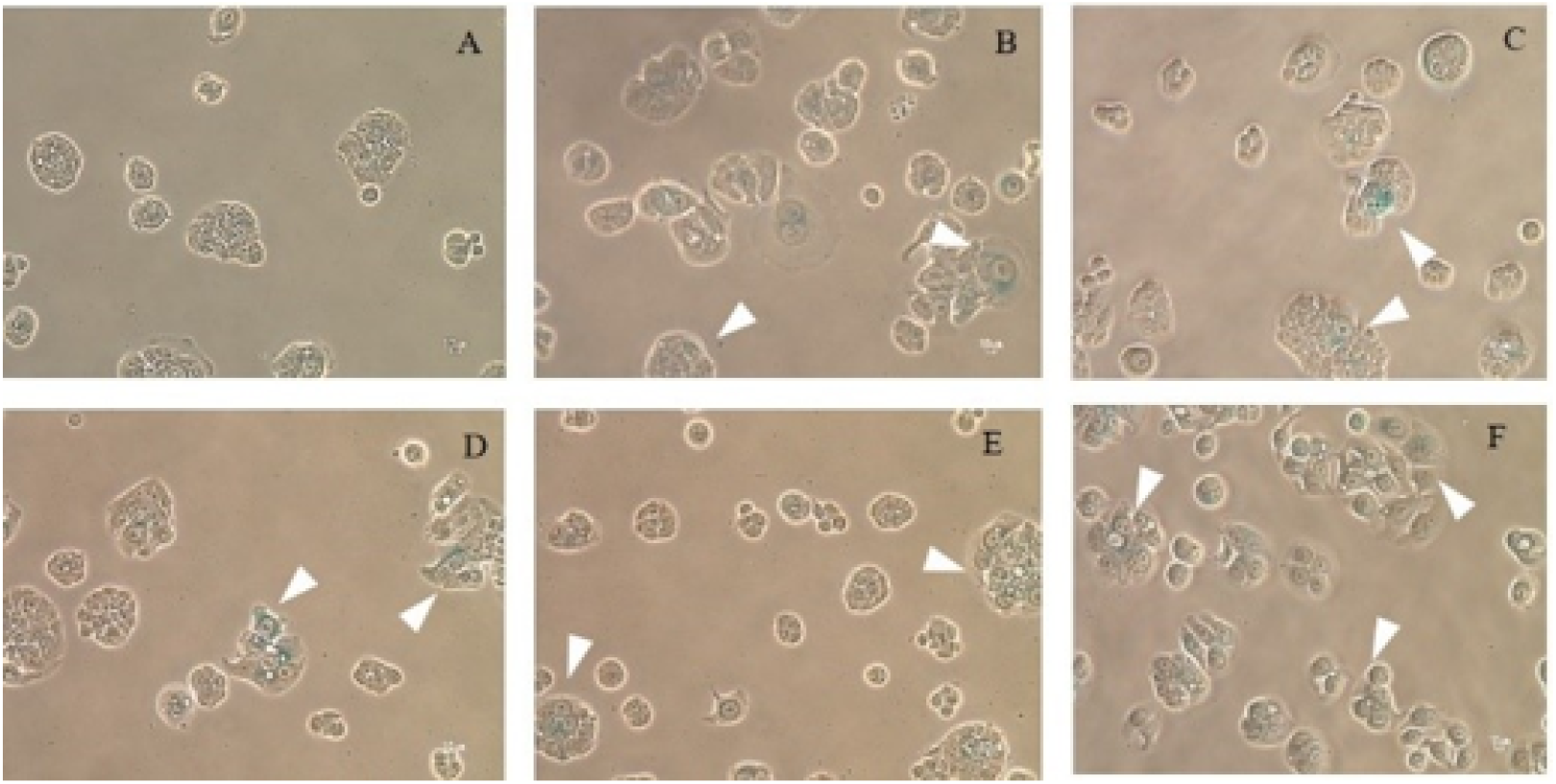
Cellular senescence inducing effect of HRBE on HT-29 cells. After treated with 0 mg/ml (A), 0.5 mg/ml (B), 1 mg/ml (C), 2.5 mg/ml (D), and 5 mg/ml (E) for 24 h compared to 0.05 mg/ml mitomycin C treatment (F). The blue stained cells with flatten and enlarged cell morphology represented the cellular senescence status indicated by the white arrow heads.

**Fig 5.**
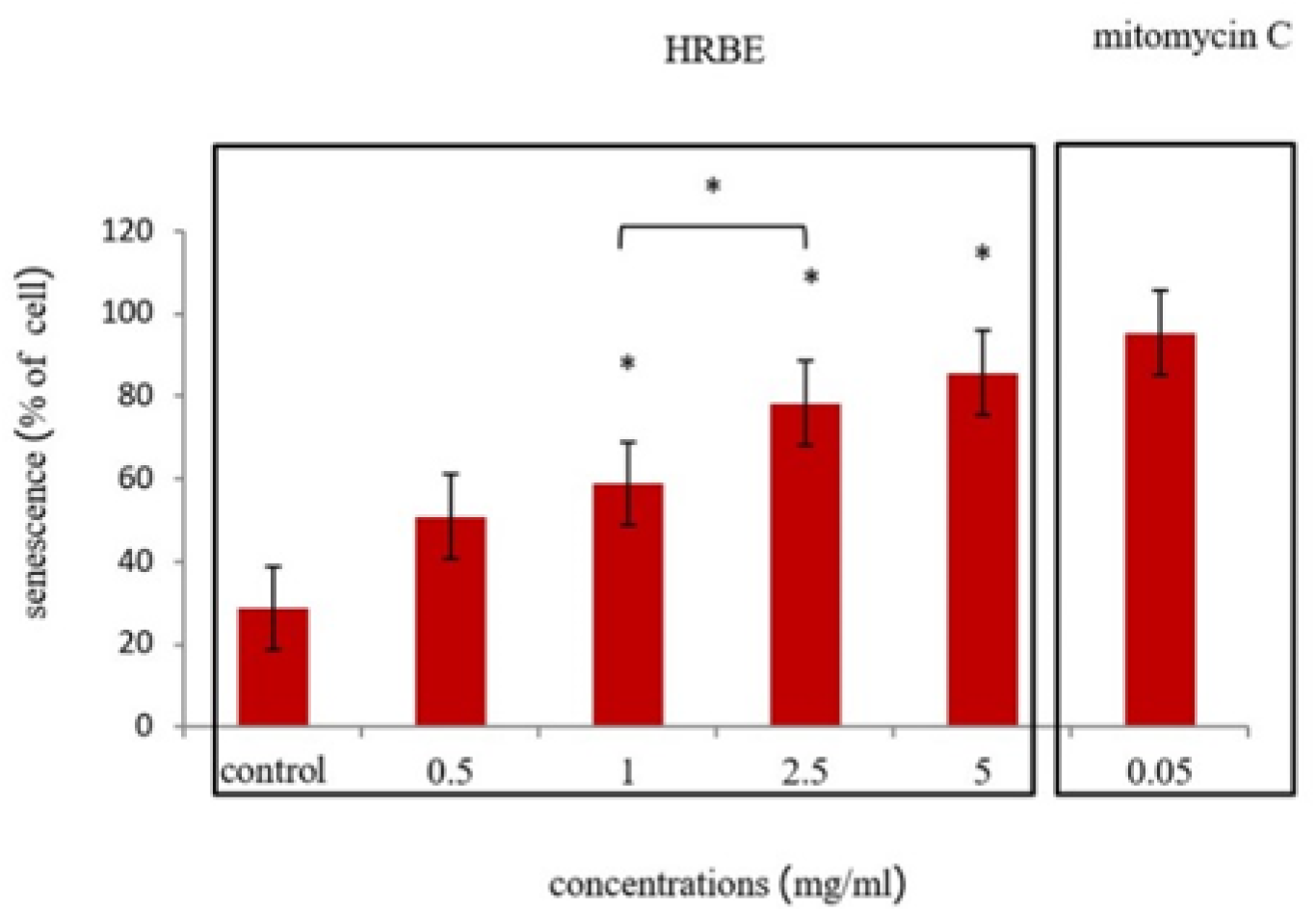
Cellular senescence inducing effect of HRBE and mitomycin C on HT-29 cells by SA-β-gal staining technique. Data represented in the mean ± SD. Statistical analysis was per-formed using student’s t – test compare each concentration with the control group (0 mg/ml) (*, p ≤ 0.05; **, p ≤ 0.01).

**Fig 6.**
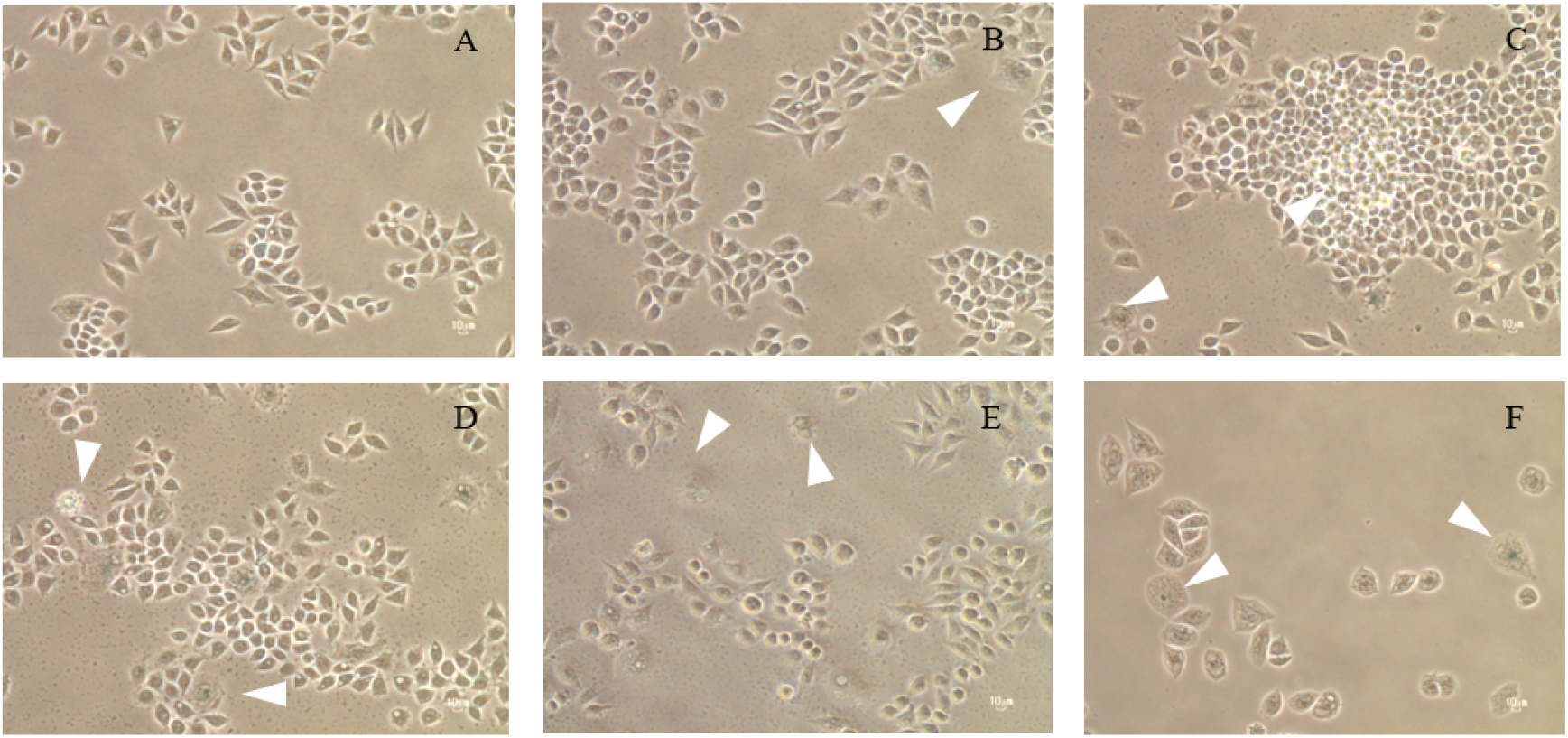
Cellular senescence inducing effect of HRBE on SW-620 cells. After treated with 0 mg/ml (A), 0.5 mg/ml (B), 1 mg/ml (C), 2.5 mg/ml (D), and 5 mg/ml (E) for 24 h compared to 0.05 mg/ml mitomycin C treatment (F). The blue stained cells with flatten and enlarged cell morphology represented the cellular senescence status indicated by the white arrow heads.

From the result in HT-29 cell line, the mean values of the stained cells were 28.65, 50.94, 58.59, 78.43, 85.74 percent of HRBE and 95.45 percent of mitomycin C (Fig 5), respectively. The SA – β – gal color was gradually increased as the concentration of the extract increased. As with the extract test with SW-620 cell line, the mean values of dyed cells were 3.75, 6.10, 16.32, 8.14, 32.76 of HRBE and 12.61 percent of mitomycin C (Fig 7). Cell dye increased with the extract concentration and HRBE can induce HT-29 to become cell senescence than SW-620.

**Fig7.**
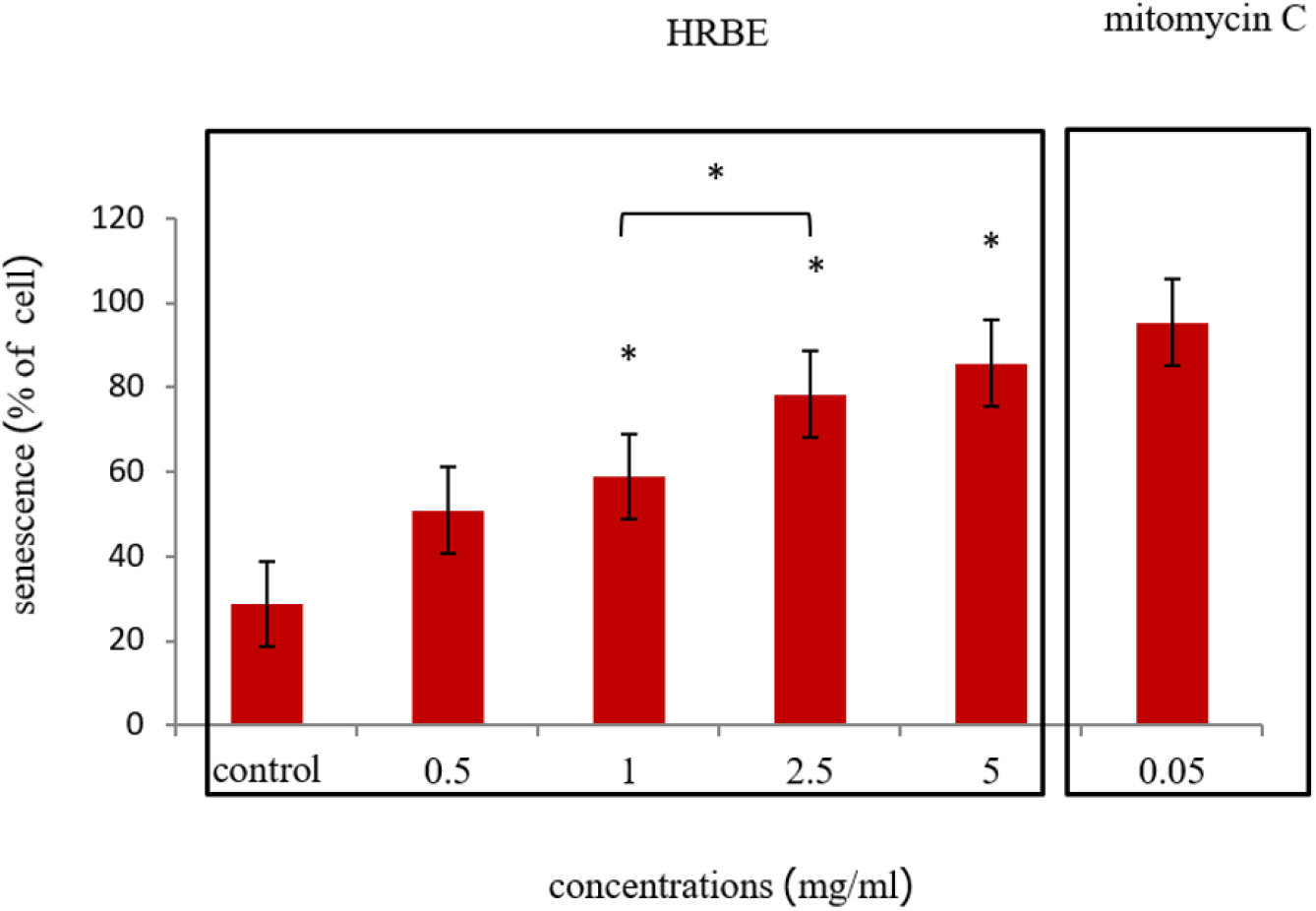
Cellular senescence inducing effect of HRBE and mitomycin C on SW-620 cells by SA-β-gal stunning technique. Data represented in the mean ± SD. Statistical analysis was per-formed using student’s t – test compare each concentration with the control group (0 mg/ml) (*, p ≤ 0.05; **, p ≤ 0.01).

### 4. Quantification of apoptosis using fluorescence microscope AO/PI double staining test

AO is a membrane permeable nuclear dye that stains both live and dead cells (early apoptosis) in green color, while PI is a membrane impermeable nuclear dye that stains late apoptosis dead cells red [9, 10]. Use for confirmed paten cell apoptosis. In result, HRBE was examined for apoptotic-inducing activity by fluorescence microscopy analysis. Morphology of SW-620 changes after treatment with HRBE at 72 h., characterized by apoptosis such as cytoplasmic shrinkage and membrane blabbing, chromatin condensation in early apoptosis and late apoptosis can indicated by red color from PI (Fig 8).

**Fig 8.**
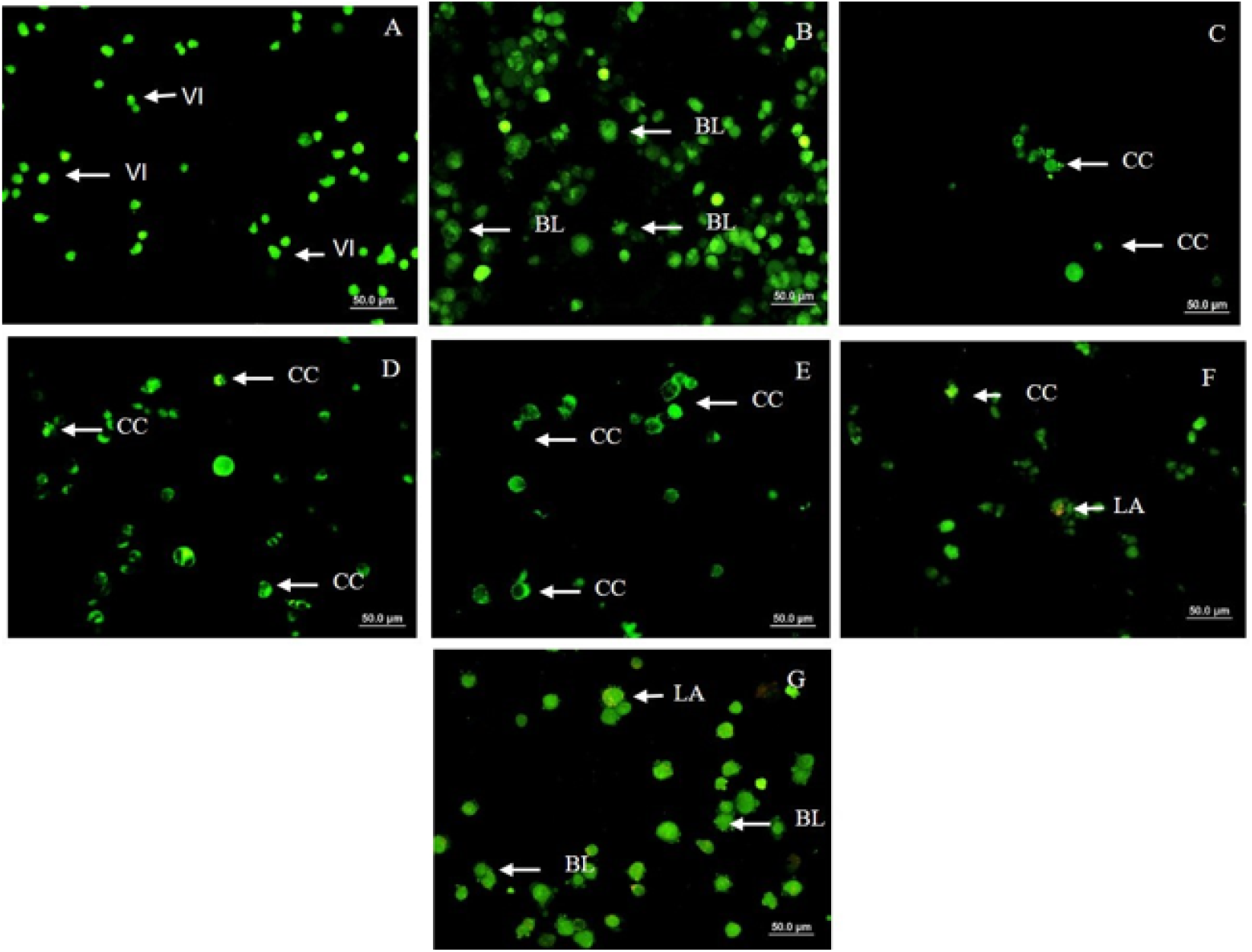
fluorescence microscope AO/PI double stained of HRBE on SW-620 cells. After treated with 0 mg/ml (A), 1.25 mg/ml (B), 2.5 mg/ml (C), 5 mg/ml (D), and 10 mg/ml (E) for 72 h compared to 0.05 mg/ml mitomycin C treatment (F) and Doxorubicin 1 μM (G). VI: Viable cell; Blabbing of the cell membrane; CC: Chromatin condensation; LA: Late apoptosis.

### 5. The analysis of cell apoptosis by Annexin V and flow cytometry

In apoptotic cells, phosphatidylserine (PS) phospholipid is flipped from the inner to the outer leaflet of the plasma membrane to signal for nearby phagocytic cells. Annexin V is a calcium dependent phospholipid-binding protein with a high affinity for PS [11]. This Annexin V staining is commonly co-stained with propidium iodide (PI) for identification the membrane potential to differentiate between early and late apoptotic cells by Flow cytometry analysis. In early apoptosis (Annexin-V positive, PI negative), late apoptosis (Annexin-V and PI positive) and necrotic (Annexin-V nega-tive, PI positive) cells [12]. Which corresponds to the results of our studies after treat-ment with HRBE at 72 h on SW-620. The data show HRBE can induce cell apoptosis in dose dependent manner and significantly when treated HRBE concentration 5 and 10 mg/ml (5mg/m/ = 8.9 ± 3%, 10mg/ml = 10.4 ± 2.3%) (Fig 9).

**Fig 9.**
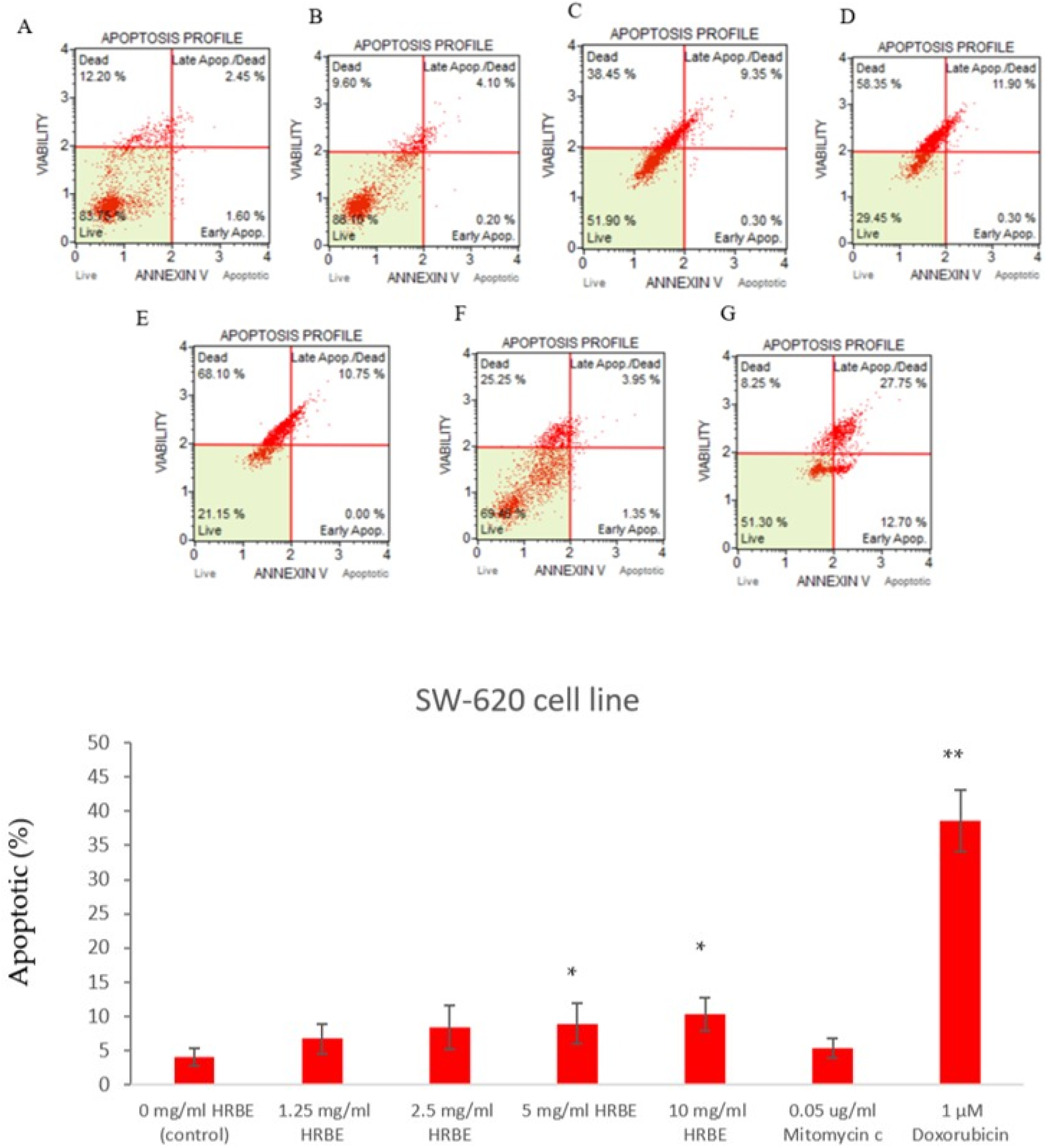
The percentage of cell apoptosis on SW-620 cells at 72h indicated by Annexin V and flow cytometry. After treated with 0 mg/ml (A), 1.25 mg/ml (B), 2.5 mg/ml (C), 5 mg/ml (D), and 10 mg/ml (E) for 72 h compared to 0.05 mg/ml mitomycin C (F) and Doxorubicin 1 μM (G). Data represented in the mean ± SD. Statistical analysis was performed using student’s t – test compare each concentration with the control group (0 mg/ml) (*, p ≤ 0.05; **, p ≤ 0.01)

### 6. The analysis of Cell cycle by flow cytometer

Cell cycle analysis used propidium iodide (PI) and RNAse in proprietary formulation that for quantitative measurement of the percentage of cell in the G0/G1, S, G2/M phase of cell cycle. In result, HT-29 cell line after treated with HRBE significantly when used concentration 5 and 10 mg/ml compared to the control. The percent number of cells were increase in G0/G1 phase (5mg/m/ = 39.63 ± 11.83%, 10mg/ml = 61.67 ± 3.76%) decrease in S phase (5mg/m/ = 22.83 ± 4.57%, 10mg/ml = 17.07 ± 2.37%), G2/M phase (5mg/m/ = 13.23 ± 2.81%, 10mg/ml = 9.5 ± 1.15%) (Fig 10).

**Fig 10.**
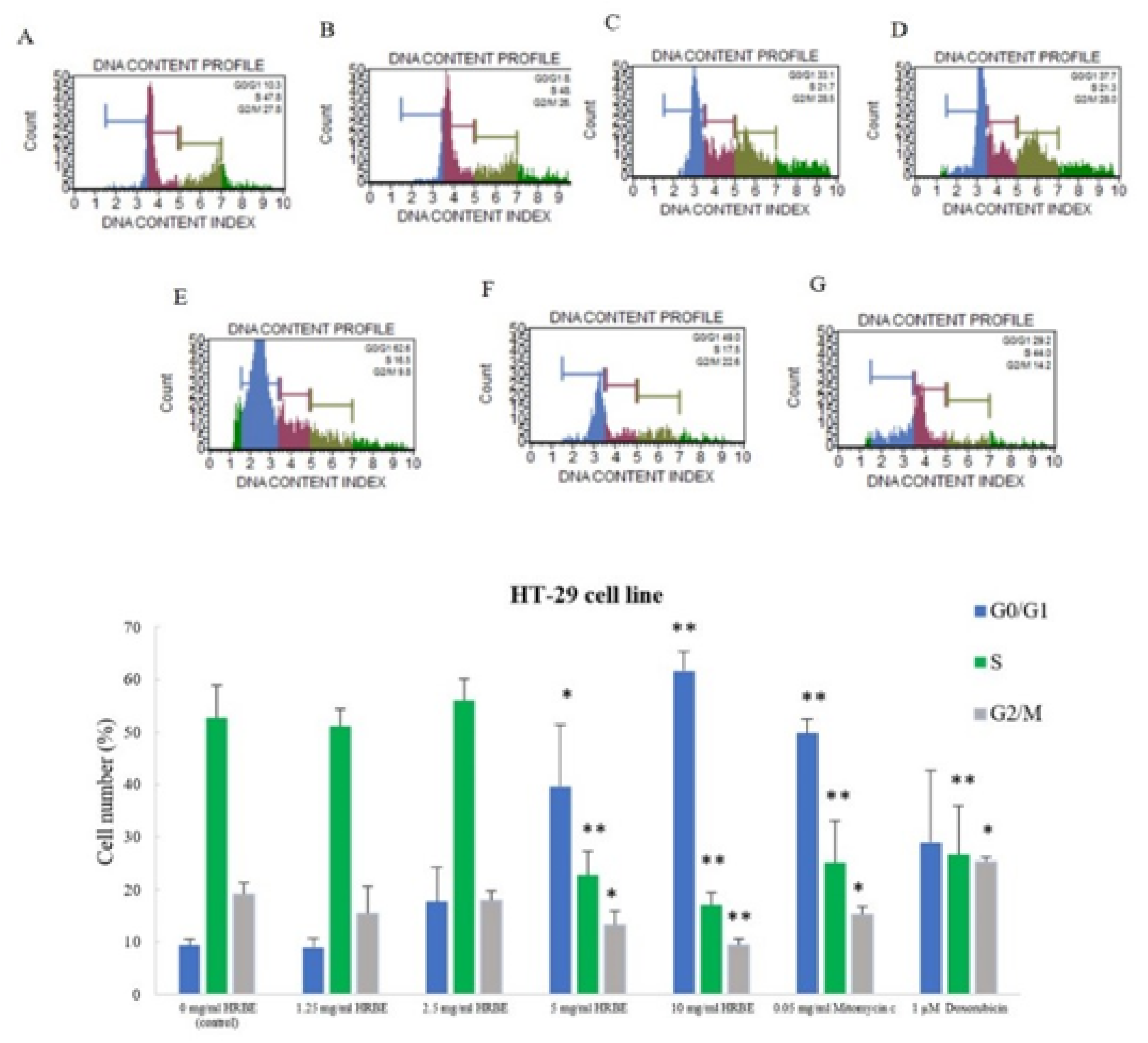
The cell cycle analysis by flow cytometer of HT-29 cells at 24h. After treatment with HRBE concentration 0 mg/ml (A), 1.25 mg/ml (B), 2.5 mg/ml (C), 5 mg/ml (D), and 10 mg/ml (E) compared to 0.05 mg/ml mitomycin C (F) and Doxorubicin 1 μM (G). Data represented in the mean ± SD. Statistical analysis was performed using student’s t - test compare each concentration with the control group (0 mg/ml) (*, p ≤ 0.05; **, p ≤ 0.01 significant).

### 7. Cytotoxicity test of fractions HRBE by MTT assay

Crude of hydrolyzed riceberry rice bran extract (HRBE) was prepared by alkaline endo-peptidase that can inhibit cancer cell growth. Therefore, we would like to isolat-ed size of peptide from crude extract can anti-cancer by centrifugal filter devices into >50, 50-30, 30-10, 10-3 kDa, and <3 kDa fractions. For confidence, the efficacy of an-ti-cancer cell growth comes from peptide. Thus, remove anthocyanins first that flavonoid compounds and produce plant colors such as blue, red and purple [13].

The result show percentage yield of fractions HRBE (Table 2) and Cytotoxicity test of fractions HRBE by MTT assay used different size of peptides with concentra-tions 0 – 10,000 μg/ml, Doxorubicin concentration 0 – 1 μM and mitomycin C concentrations 0 – 100 μg/ml for 24 h, 48 h and 72 h. on SW-620 (Fig 11). The lower of IC_50_to indicates the growth inhibitory activity for SW-620 of rice-bran peptide is present in the >50 kDa fraction. Moreover, IC_50_ of the >50 kDa fraction (4,908 μg/ml) that less than the IC_50_ of crude extract (5,492 μg/ml) at 72 h. (Table 3)

**Table 1.**
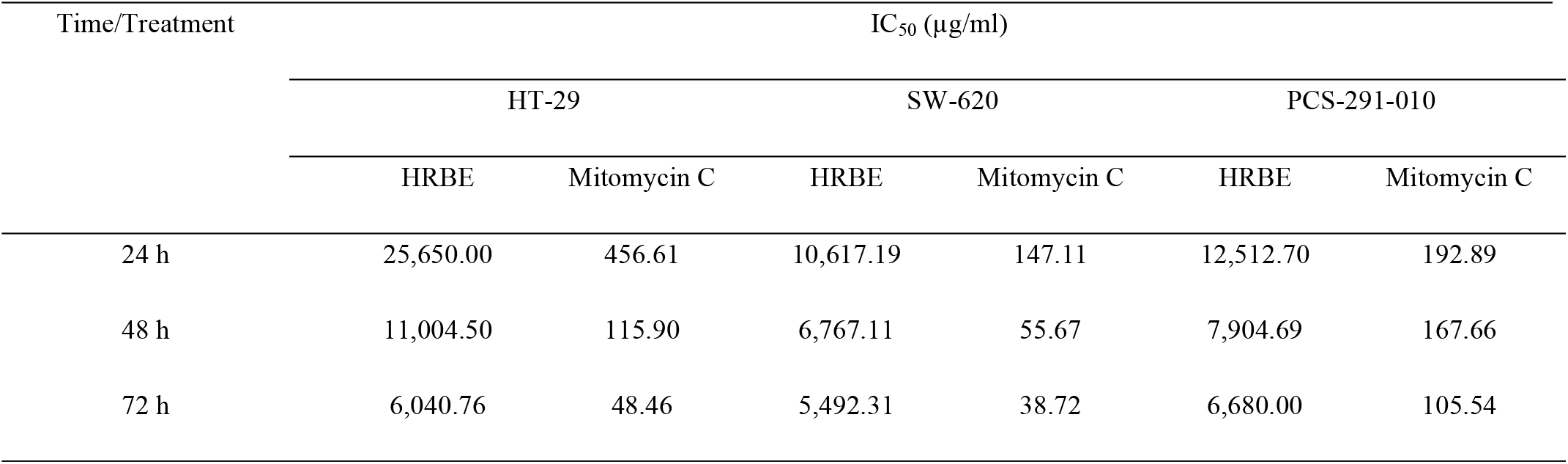
IC_50_ values of HRBE and mitomycin C on tree different cell lines.

| Time/Treatment | IC<sub>50</sub> (μg/ml) |  |  |  |  |  |
| --- | --- | --- | --- | --- | --- | --- |
|  | HT-29 |  | SW-620 |  | PCS-291-010 |  |
|  | HRBE | Mitomycin C | HRBE | Mitomycin C | HRBE | Mitomycin C |
| 24 h | 25,650.00 | 456.61 | 10,617.19 | 147.11 | 12,512.70 | 192.89 |
| 48 h | 11,004.50 | 115.90 | 6,767.11 | 55.67 | 7,904.69 | 167.66 |
| 72 h | 6,040.76 | 48.46 | 5,492.31 | 38.72 | 6,680.00 | 105.54 |

**Table 2.**
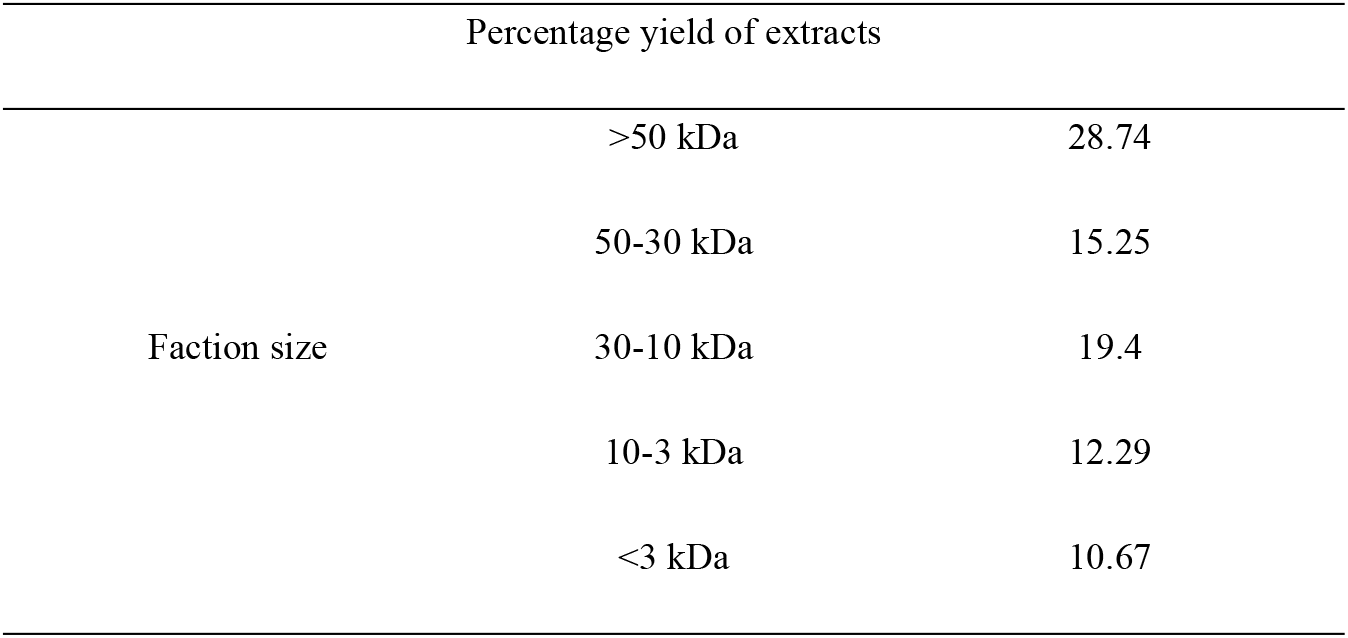
Percentage yield of faction HRBE.

**Table 3.** IC_50_ values of faction HRBE, doxorubicin and mitomycin C for 24 h, 48h and 72 h.

| Time/Treatment |  | IC<sub>50</sub> |  |  |  |  |  |  |
| --- | --- | --- | --- | --- | --- | --- | --- | --- |
|  |  | Faction of HRBE (μg/ml) |  |  |  |  | Doxorubicin | Mitomycin C |
|  |  |  |  |  |  |  | (μM) | (μg/ml) |
|  |  | >50 kDa | 50-30 kDa | 30-10 kDa | 10-3 kDa | <3 kDa |  |  |
| 24h |  | 471,623.00 | -28923.60 | -29630.01 | 103,029.70 | 292,099.90 | 0.50 | 33.34 |
| 48h |  | 10,837.16 | 26,148.02 | 22,442.69 | 23,037.82 | 18,488.71 | 0.08 | 12.74 |
| 72h |  | 4,908.97 | 8,519.21 | 11,602.07 | 13,428.92 | 11,116.36 | 0.06 | -1.79 |

**Fig 11.**
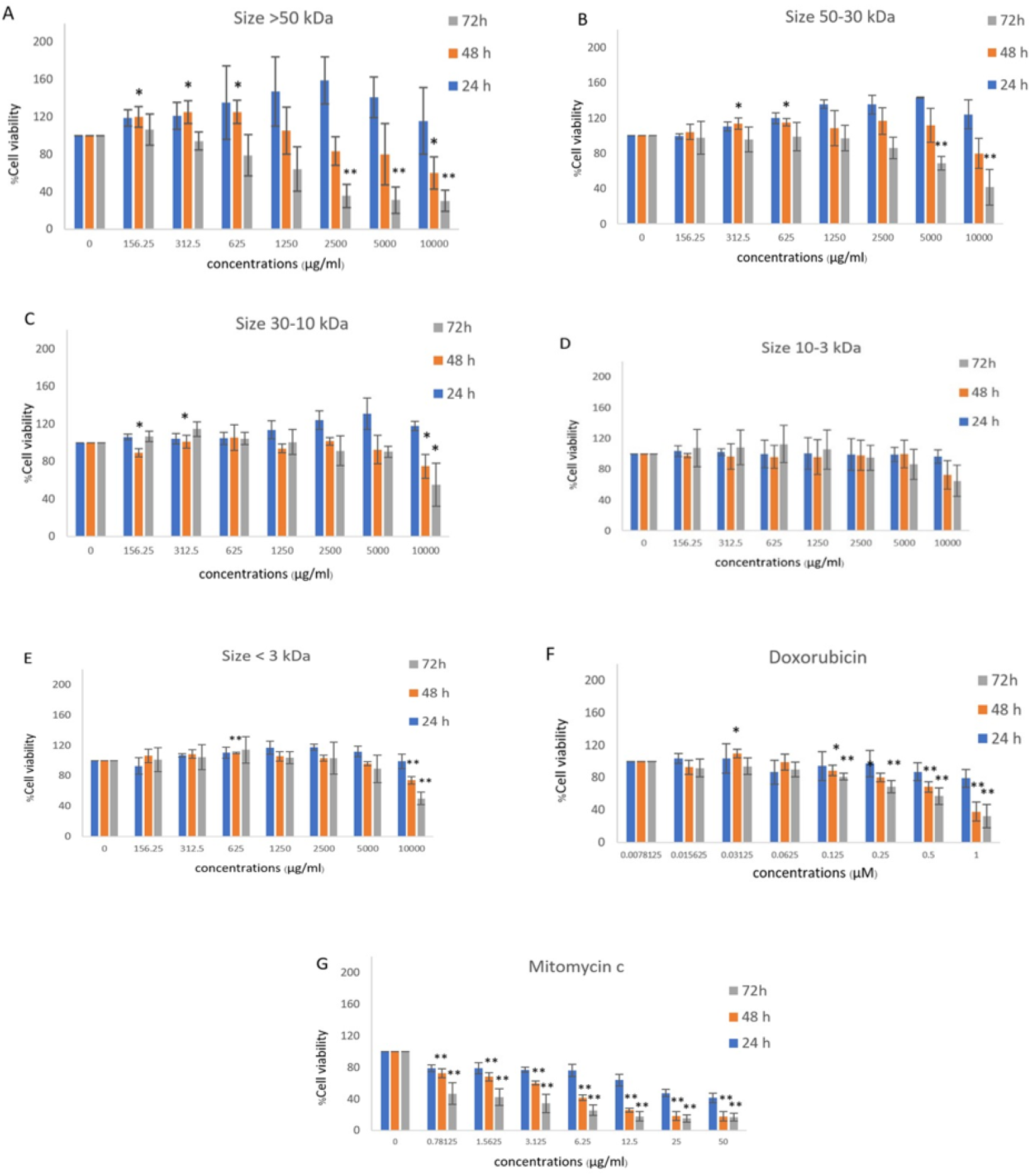
Cytotoxicity test by MTT assay on SW-620 cells after treated with faction of HRBE. Concentrations 0 – 10,000 μg/ml including (A)>50, (B)50-30, (C)30-10, (D)10-3 kDa and (E)<3 kDa fractions, (F)Doxorubicin concentration 0-1 μM and (G)mitomycin C concentrations 0 – 50 μg/ml for 24 h, 48 h and 72 h. The percentage of cell viability was determined as compared to an untreated condition. Data represented in the mean ± SD. Statistical analysis was per-formed using student’s t – test (*, p ≤ 0.05 significant; **, p ≤ 0.01 significant.

### 8. Quantification of apoptosis using fluorescence microscope AO/PI double staining test with >50 kDa fraction of HRBE

For confirming >50 kDa fraction HRBE can induce SW-620 in apoptosis at 72h. The morphology of SW-620 after treated with different concentration of >50 kDa fraction HRBE showed characterize cell apoptosis including membrane blabbing, chromatin condensation in early apoptosis and late apoptosis can be indicated by red color from PI (Fig 12) This result indicates that the >50 kDa fraction of the peptide in HRBE was effective to inhibit cancer cells by inducing apoptosis.

**Fig12.**
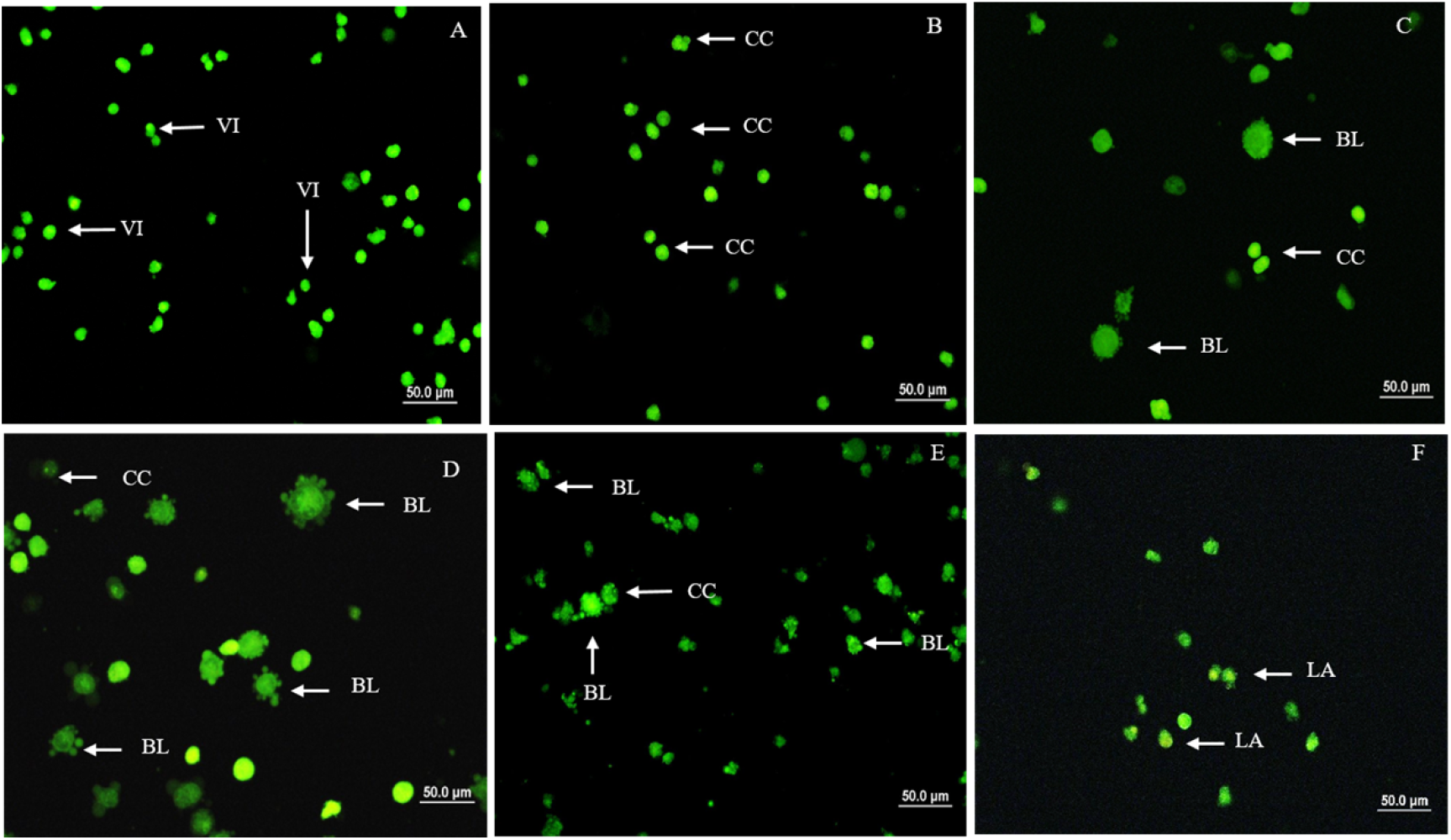
fluorescence microscope AO/PI double stained of fractions HRBE on SW-620 cells. After treated with 0 mg/ml (A), 2.5 mg/ml (B), 5 mg/ml (C) and 10 mg/ml (D) for 72 h compared to 0.05 mg/ml mitomycin C treatment (E) and Doxorubicin 1 μM (F). VI: Viable cell; Blabbing of the cell membrane; CC: Chromatin condensation; LA: Late apoptosis

## Discussion

Colon cancer is the third most common cancer and the second leading cause of cancer related death worldwide [14]. There is a study of natural substances to replace or use in conjunction with chemotherapy drugs. And it has been reported that proteins from nutrients more efficient release of bioactive peptides and selective anti-cancer. In the present study, cytotoxicity test on fibroblast normal human cell line (HDFa), non – metastasis human colon cancer cell line (HT-29) and metastasis human colon cancer cell line (SW-620) with hydrolyzed riceberry rice bran extract (HRBE) by MTT assay. The percentage of cell viability after treatment decreased when increasing the concen-tration of extract (dose-dependent) and considering the IC_50_ was effective when treated HRBE at 48 and 72 h (time-dependent). A similar observation study of Banjerd-pongchai et al. (2009) tested the cytotoxicity of purple glutinous rice extract on human hepatocellular carcinoma (HepG2) and prostate cancer (LNCaP) were found effective at 48 h were better than 24 h. The efficacy of rice bran extract on colon cancer cells (HCT-116) and breast cancer (HTB-26) showed that the cytotoxicity rate of both cancer cells increased dependent on the time [15]. The comparison IC_50_ of three cell lines when treated HRBE and mitomycin C for positive control that show the IC_50_ of HT-29 were higher than SW-620. Because the morphology of non-metastasis cancer cells and have denser adhesion between cells (cell-cell junction) than metastasis cancer that loss of cell junction and change the morphology for spread to another organ. This provides the extract affect SW-620 better than HT-29.

Therefore, from the result of MTT assay, we selected SW-620 to investigate the HRBE can induce apoptosis in the next experiment. Gel-electrophoresis technique show DNA fragmentation pattern that one of characterized in apoptosis cells. The ac-tivation of endonucleases, particularly the caspase-3 activated DNase. CAD with sub-sequent cleavage of nuclear DNA into internucleosomal fragments of roughly 180-200 base pairs (bp) [16]. Moreover, we confirmed paten cell apoptosis in morphology by fluorescence microscope AO/PI double staining test. The morphological apoptosis cell such as membrane blabbing and chromatin condensation in early apoptosis. And late apoptosis can indicate in red color from PI that binds to DNA but cannot passively traverse into cells that possess an intact plasma membrane [9, 10] because the cell membrane of normal cell and cancer cells were different. In normal cells have zwitterion phosphatidylcholine and sphingomyelin in an outer leaflet and anionic phosphatidylserine and the phosphatidylethanolamine in the inner leaflet with the asymmetric distribution. But in the cancer cell was loss asymmetric distribution, in-creased negative charges and alteration PH on the cell membrane [17]. This is a cause why PI can indicate late apoptosis including the efficacy of the extract on cancer cells and cause of the difference in the activity of the HRBE between HT-29 (non – metasta-sis) and SW-620 (metastasis) for the quantity of peptide to enter cells.

In apoptotic cells, phosphatidylserine phospholipid is flipped from the inner to the outer leaflet of the plasma membrane to signal for nearby phagocytic cells. Annexin V is a calcium dependent phospholipid-binding protein with a high affinity for phosphatidylserine [11]. Therefore, the percentage of apoptosis on SW-620 after treated with HRBE can detect by Annexin V and flow cytometry. Our result shows that HRBE at concentrations 5 and 10 mg/ml were significantly increased to induce cell apoptosis. Which is consistent with previous studies in other rice such as the water extract of brewers’ rice (WBR) on HT-29 show characteristics of apoptosis including early and late apoptosis at 48h [18] and induce apoptosis by the significant activation of caspase-3 and −8 activities [19]. In Purple rice extract by methanol, cultivar exhibited the highest inhibitory growth effect on human hepatocellular carcinoma cells and abil-ity to induce apoptosis via the mitochondrial pathway with the loss of mitochondrial transmembrane potential (MTP) and activation of caspase-3 and −9 [20]. Moreover, ex-traction of chemical components from the Rice berry bran can induce HL-60 and Ca-co-2 to apoptosis was stimulated by p53 protein that was increased to activation of caspase-3[3]. For the p53 tumour suppressor protein was regulated genes to control cell cycle in normal cells and induced cell cycle arrest for the repair of DNA, cellular senescence, and programmed cell death such as apoptosis to remove irreversibly damaged cells [21]

Therefore, an interesting study of anti-cancer was cell cycle arrest (Senescence). Cellular senescence is the stage of irreversible cell cycle arrest. Cell senescence by SA-β-gal staining show present senescence on HT-29 than SW-620 that can indicate HRBE efficiency not only induce apoptosis but can induce cell senescence. The inhibi-tion of cancer cells also depends on the type of cells. Senescence cells clued be initially undergone cell cycle arrest that decreased number of cells in S, M/G2 phase and in-creased number of cells in G1/G0 phase. For flow cytometer, can quantitative meas-urement of the percentage of cell by propidium iodide (PI) and RNAse. PI was specific DNA stunning that can indicate the different stage of cell cycle because in the G1 phase have constant DNA content (1 x), cells in the G2 /M phase have constant DNA (2 x) twice the intensity of the G1 phase [22]. This study demonstrate HRBE can increase the number of cells in the G0/G1 phase and decrease in S, G2 and M phase that indi-cates HRBE can induce HT-29 cell line to senescence. Because cyclin-dependent ki-nase(CDKs) and cyclin are regularly protein control role of cell cycle and CDK-cyclin complexes can be regulated by a group of proteins called cyclin-dependent kinase in-hibitors (CDKIs) such as p21that is downregulation of p53. The p21 can be inducing G1 cell cycle arrest by inhibiting the activity of cyclinCDK2/4 complexes, E2F, and PCNA [23]. Rice berry bran extract by dichloromethane on HL-60, MCF-7 and Caco-2 were decreasing Cyclin B1 that complex with Cdk1 control S to G2 phase in the cell cycle [3]. Moreover, the HT-29 was treated with Brewers’ rice at 48h. was an increase in the cell population at the sub-G0 phase [18].

Then we identified the specific size of protein in the crude extract to anti-cancer by centrifugal filter devices into >50, 50-30, 30-10, 10-3 kDa, and <3 kDa fractions and investigate cell viability on SW-620 cell line to determine the size of proteins that in-duce apoptosis. The lower IC_50_ can indicates the growth inhibitory activity of rice-bran peptide was present in the >50 kDa fraction. Moreover, IC_50_ of >50 kDa fraction (4,908 μg/ml) less than the IC_50_ of crude extract (5492.31 μg/ml) at 72 h. This indicates that Peptides of >50 kDa fraction were effective in inhibiting cancer as well. And >50 kDa fraction that shows characterized by apoptosis such as membrane blabbing and chro-matin condensation when used fluorescence microscope AO/PI double staining test. Rice bran was a rich source of dietary fibres and proteins such as albumin, globulin, glutelin and prolamin. Albumins were separated into three to four sub-fractions using gel filtration chromatography and have 100 kDa or less molecular weight when ana-lyzed by size-exclusion HPL [24]. According to a previous study, Alcalase is an alkaline endo-peptidase that gave higher oil and protein yields. Its activity in hydrolyzing glu-telin, as the main protein accounting for 60–80% of total protein in rice, was higher than neutral protease. Glutelin was soluble in alkaline solution and hence easily cleaved by alkaline peptidase [7, 8]. Glutelin (MW 57 kDa) as the main protein in rice was mostly hydrolyzed by alcalase [25].

## Conclusions

In conclusion, the results show that the efficacy of HBRE can inhibit colon cancer cell line depends on time, dose and cell type including inducing to apoptosis in SW-620 and senescence in HT-29 that decreased the number of cells in S, M/G2 phase and increased number of cells in G0/G1 phase. Moreover, for >50 kDa fraction of HRBE demonstrate the ability to induce apoptosis and cytotoxicity better than the crude version. This finding would be useful and applicable for medical research and colon cancer treatment in the future.

## Acknowledgements

This research was supported in part by the Graduate Program Scholarship from the graduate school and Department of Zoology, Faculty of Science, Kasetsart University.

## Author Contributions

Conceptualization, P.C. and M.K.; methodology, P.C., M.K. and V.W.; validation, P.C. and V.W.; investigation, P.C.; resources, S.T., P.C. and M.K.; writing—original draft preparation, V.W.; writing—review and editing, P.C.; supervision, P.C. and M.K.; project administration, P.C.; funding acquisition, P.C. and M.K. All authors have read and agreed to the published version of the manuscript.

